# Democratizing water monitoring: Implementation of a community-based qPCR monitoring program for recreational water hazards

**DOI:** 10.1101/2020.02.13.947259

**Authors:** Sydney P Rudko, Ronald R Reimink, Bradley Peter, Jay White, Patrick C Hanington

**Author notes:** Corresponding author: Patrick Hanington.

## Abstract

Recreational water monitoring can be challenging due to the highly variable nature of pathogens and indicator concentrations, the myriad of potential biological hazards to measure for, and numerous access points, both official and unofficial, that are used for recreation. The aim of this study was to develop, deploy, and assess the effectiveness of a quantitative polymerase chain reaction (qPCR) community-based monitoring (CBM) program for the assessment of bacterial and parasitic hazards in recreational water. This study developed methodologies for performing qPCR ‘in the field’, then engaged with water management and monitoring groups, and tested the method in a real-world implementation study to evaluate the accuracy of CBM using qPCR both quantitatively and qualitatively. This study found high reproducibility between qPCR results performed by non-expert field users and expert laboratory results, suggesting that qPCR as a methodology could be amenable to a CBM program.

## 1.0 INTRODUCTION

Community based monitoring is now routinely used for conservation and environmental monitoring(1). Citizen science describes both a methodology of conducting large-scale research by recruiting volunteers, and refers to the process by which citizens are involved in scientific investigation as researchers. Citizen science can include community based monitoring (CBM) as a process of collaboration between government, industry, academia, and local community groups to monitor, track, and respond to issues (2–4).

The earliest incarnations of citizen science and CBM relied on volunteers as data collectors, but the discipline of CBM has grown and evolved. Recent arguments in favor of CBM suggest the field move away from a paradigm of “using citizens to do science” to an equal power relationship which views citizens as scientists, embracing some of the ideals of participatory action research (5).

CBM is poised to improve environmental decision-making. Its use has been on the rise due to budgetary constraints in both government and academia, but also because CBM can be a powerful methodology for generating large spatial or temporal datasets for monitoring/surveillance purposes. CBM improves scientific literacy, builds social capital, improves participation in local issues and benefits the environment (6,7). Traditional CBM programs have typically relied on volunteers to conduct biodiversity surveys, to conduct simple tests (i.e. Secchi disk tests for assessing water clarity), or to collect specimens and send them to central facilities for analysis. However, modern monitoring methods conducted in academia, industry, and government have evolved considerably to include large-scale spatial assessment methods, for example: algal/cyanobacteria bloom-tracking satellites, next generation sequencing analysis, and eDNA monitoring. CBM programs also must evolve and advance as new technologies become available. In water monitoring especially, quantitative polymerase chain reaction (qPCR) has emerged as the method of choice for conducting routine compliance monitoring of water bodies (8).

Quantitative PCR methods for the detection of surrogates and hazards in water have existed for decades and can be used to detect minute quantities of an organisms’ DNA in a complex matrix such as water, soil, or blood. qPCR is highly sensitive (in theory, capable of detecting a single copy of organismal DNA) and is very specific for particular regions of DNA. In the last decade, agencies responsible for monitoring the environment and health have begun to capitalize on the potential of qPCR. Some of the greatest strides have been made in health, especially after the USEPA EMPACT study, which found that levels of enterococcus as measured by qPCR correlate with risk of human gastrointestinal illness (9). Since then, strides have been made in correlating the amount of human-associated *Bacteroides* with human health targets (10,11). Screening for toxigenic cyanobacteria species is also moving towards molecular detection method. For example, in Poland, initial screening for toxin genes in recreational waters is conducted using qPCR, followed by immunochemical analysis to quantify the toxins (12). In related fields like environmental monitoring some locales have moved to molecular methods for monitoring for the veliger stage of invasive zebra (*Dreissena polymorpha*) and quagga (*Dreissena rostriformis bugensis*) mussels.

As the effectiveness of qPCR diagnostic tests continues to be realized, it is apparent qPCR is an excellent choice for CBM, or more broadly, a decentralized monitoring system. qPCR is a platform, and with the infrastructure in place, monitoring for additional targets becomes a matter of designing/validating a new test and running it on the established infrastructure. For this reason qPCR and related molecular techniques have been touted as grand solutions for point of care diagnostics in infectious disease monitoring, this future has not yet been realized (13,14). The idea of portable diagnostic technologies that can be used to detect multiple targets, which feed information into a surveillance system, is attractive for a number of reasons, but the development to implementation gap is often wider than one would expect.

It is often presumed that highly skilled personnel are required to execute molecular biology methods such as qPCR. Additionally, technologies to conduct testing portably have only just begun to emerge onto the market and have not been fully vetted. This study is, to our knowledge, the first of its kind to test the rigor of qPCR for detection/quantification of biological hazards and their surrogates in water through a CBM-implementation study. Here, we test the feasibility, reproducibility and reliability of implementing portable qPCR water monitoring amongst a variety of groups (government, NGO, and private enterprise). This was assessed both quantitatively, by conducting our own measurements on CBM partner samples, and qualitatively, through surveying our user groups to capture their perceptions of the technology and its fit within their individual contexts and organizations.

## 2.0 MATERIALS AND METHODS

### 2.1 Implementation study design

We first connected with relevant stakeholders of recreational water in Alberta, and worked with them to determine their monitoring goals. Using a participatory research (PAR) approach, we then developed qPCR tests and testing methodologies that would fill these needs(15). Under this PAR approach, CBM partners selected study sites they felt would be appropriate, and we advised and assisted in this selection where it seemed appropriate. Since the goal of this study was to measure the effectiveness of a CBM monitoring program in a real world context, participants in the study were instructed to collect a duplicate sample or cut the filter membrane in half after filtration and send this to the university lab. Samples in our lab would be processed in an identical fashion to the field user to compare novice versus expert methodologies (Fig 1). Additionally, CBM partners sent their extracted DNA to our lab, which enabled us to also perform qPCR on their DNA extracts and to perform inhibition reactions.

**Fig 1.**
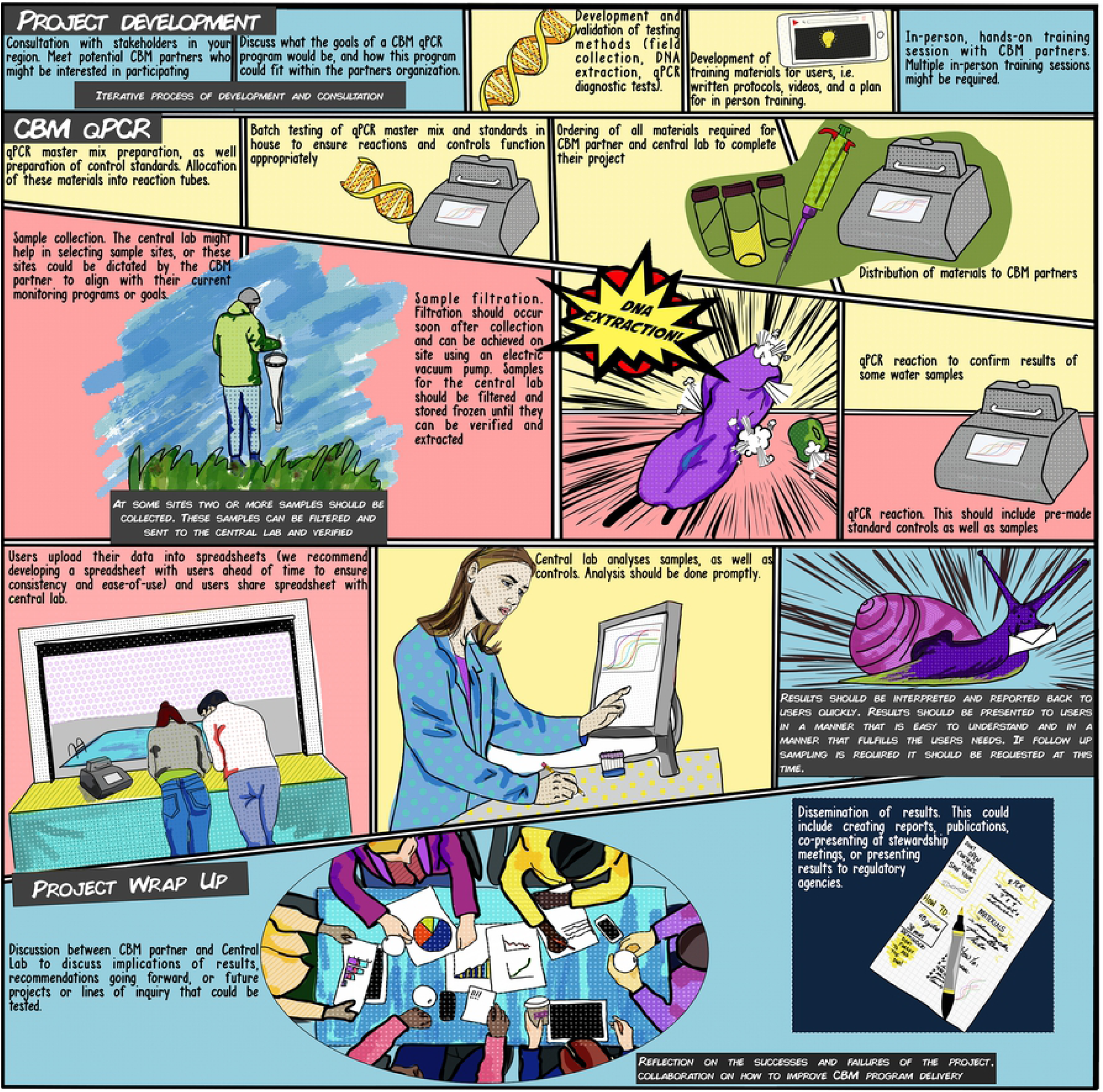
Implementation process of the CBM qPCR program. Cells with blue backgrounds are processes done in collaboration between the central laboratory and the CBM partners, yellow backgrounds indicate processes completed by the central laboratory, and red backgrounds indicate processes completed by the CBM partners.

### 2.2 Sample collection

Specific water collection methods are detailed below for each target of interest; regardless of the volume collected, all samples were then filtered through a 0.4 µm polycarbonate filter (Pall FMFNL1050) using an electric vacuum pump (Vaccubrand^®^). CBM partners had the option of either collecting and filtering a duplicate water sample for analysis, or cutting their filter membranes in half to be analyzed at the university lab.

#### 2.2.1 Avian schistosome monitoring

Sample collection was conducted as described in Rudko et al (2018). Briefly, 25L water samples, collected one litre at a time across a shoreline up to ∼1m deep were passed through a 20µm plankton tow. Debris from inside were washed down using well water (this is not a contamination risk when monitoring for avian schistosomes as these parasites are shed from snail hosts, and only when those snail hosts also co-occur with locations where the bird definitive host’s feces are also present(16)) followed by a 95% ethanol wash and collection in sterile 50-mL conical tubes.

#### 2.2.2 Toxin-producing cyanobacteria monitoring

Sample collection was conducted from watercraft operated by CBM partners on various lakes. Samples were collected through a one-way foot valve attached to weighted 3/4” Nalgene tubing. Samples were only collected from the euphotic zone as determined by a Secchi disk measurement at each lake’s deepest point. Ten sampling locations were selected for each lake, with water being composted from each sampling location into a central container. Water from this container was then poured into 50-mL conical tubes. Equipment was decontaminated between lakes using quaternary ammonium compound to prevent contamination between lakes.

#### 2.2.3 HF183 monitoring

All samples were collected by scooping two 50ml samples in sterile, conical, collection tubes from the surface water 15m from shore every 150m along the entire perimeter of each participating lake.

### 2.3 DNA extraction

#### 2.3.1 In field method

DNA extraction was conducted using the MI Sample Prep Kit (Biomeme) according to the manufacturers’ instructions. The MI sample prep kit is designed to function in the field. Lysis is accomplished by placing a filter in the lysis buffer and shaking for one minute. Next, the solution is passed through a syringe unit fitted with a DNA binding column. The column undergoes two washes to remove proteins and salts, and then is dried using an acetone buffer before elution. In 2018, the avian schistosomes monitoring group was interested in transitioning to a DNA extraction method that would allow for batch processing of samples. We therefore opted to transition their program to the DNAeasy DNA extraction kit (Rudko et al. 2018). To set up this remote laboratory in a cost-effective manner, equipment (centrifuge, heating block, and vortex) were sourced from Dot Scientific (Supplementary Table 2), and pipettes were from VWR. Sample blanks were conducted by partners every batch of 24 samples processed.

### 2.4 qPCR methods

#### 2.4.1 Maintaining workflows

All master mix components were mixed in a clean room located at the University of Alberta and aliquoted into 0.2 ml thin wall PCR tubes (Axygen). All plasmid dilutions and preparation of positive controls occurred in a deadbox. Standards and reaction tubes were prepared independently to prevent cross contamination.

#### 2.4.2 In lab qPCR method

Samples were quantitated relative to a plasmid standard curve which contained 50,000, 5000, 500, 50, 5 and 0.5 copies. Each of the gene targets below was synthesized (IDTDNA) into a puc19 plasmid vector (Genscript). Thermocycling was performed on the ABI 7500 Fast or the QuantStudio 3 using a standard, 40 cycle, two-step reaction. The thermocycling parameters were a 30 second hold at 95 degrees, followed by a 30 second denaturation cycle at 95 degrees, and a 60 degrees annealing cycle. Each qPCR reaction had a final volume of 20, and we added 5μL of DNA to each reaction.

#### 2.4.3. Avian schistosomes

The *18S* avian schistosomes-targeting qPCR assay was performed as described in Narayaran et al (2015) and Rudko et al (2018). The LOD_95_ of this technique is 3.4 gene copies/rxn (17) (Table 1S). qPCR master mix (IDT DNA) containing 1x Master mix, and 200nm forward reverse primer and fluorescein-labeled probe was used.

#### 2.4.4 Toxigenic (mcyE gene) cyanobacteria monitoring

The mcyE gene targeting qPCR assay was performed as described in Qiu et al. (2013) and Sipari et al. (2010) (Table 1S). The LOD_95_ of this technique is 6.25 copies/5μL. qPCR master mix (IDT DNA), containing 1x Master mix, and 200nm forward reverse primer and 125nm fluorescein-labeled probe was used.

#### 2.4.5 HF183 bacteroides monitoring

This 16S gene-targeting assay was performed as described in Haugland et al. (2010). The LOD_95_ of this technique is 7.2 gene copies/rxn. qPCR master mix (IDT DNA), containing 1x Master mix, and 100nm forward reverse primer and 80nm fluorescein-labeled probe was used (Table 1S).

#### 2.4.6 In-field qPCR method

Mastermix components and concentrations were unchanged between the lab method and the field method, nor were the thermocycling parameters. CBM partners received 4 control tubes, which consisted of a negative control, and 3 standards (5000,500, and 50 copies). They were instructed not to open these tubes to prevent contamination. CBM partners also received 12 tubes to add their own samples DNA to (Fig 1).

#### 2.4.7 Inhibition controls

Inhibition controls were performed as described in Rudko et al. (2017) (20). Plasmid control DNA was spiked in excess into qPCR reactions containing 5μL of water sample DNA, and inhibition was defined as a 3-ct (i.e. 1 log) shift in amplification.

### 2.5 Creation of the field kits

Field kits given to CBM partners contained: the M1 DNA extraction kit (Biomeme), 1.5 ml snap-cap tubes, sample collection vials (Corning), a 20 micron plankton tow (Acquatic Research Instruments), 0.45 uM polycarbonate filter funnels (Pall, FMFNL 1050), a 20 μL pipette, a box of pipette tips, PCR tubes, a laptop (some Acer, some Chromebook) an Open qPCR thermocycler, all the necessary cables, and reaction strips (Fig 1, Supplementary Materials).

### 2.6 Training of CBM partners

CBM partners were provided with a training video, and a written protocol. Additionally, they were provided with two in-person training session. Typically we would demonstrate the method in our laboratory, and the second training session would on-site at their location, where the CBM partner would run their first samples.

### 2.7 Capturing CBM partners perceptions of the method

CBM partners (6 in total) were administered a survey with open-ended questions regarding the implementation of the method (Supplemental Table 3). All 6 CBM partners submitted a completed survey. Surveys were blinded from the researchers to encourage honesty from participants; a research associate received the surveys via email and edited them to remove any personal identifiers before sending them to the analyst. Data were analyzed using deductive thematic analysis(21). Open coding was used, and codes were developed and modified as the analysis took place. Analyzing the codes enabled the identification of initial themes; these preliminary themes were refined to demonstrate interesting patterns in the data that were important to the successes or failures of the implementation. Themes were realized semantically (i.e. the explicit or surface meaning of the data), and latently, to identify and examine underlying ideas and assumptions that inform the semantic content of the data (22).

### 2.8 Ethics Statement

All procedures performed in studies involving human participants were in accordance with the ethical standards of the institutional and/or national research committee and with the 1964 Helsinki declaration and its later amendments or comparable ethical standards. This research was approved by the University of Alberta Human Research Ethics Board: Approval # Pro00048511.

### 2.9 Bland-Altman plots

Bland-Altman plots were created in GraphPad Prism 8 on the log transformed copy number per 5μL data. Log transformation was performed prior to conducting the analysis because this method assumes that the SD of method differences is uniform across measurements, but it has been documented that variability in measurement becomes greater when a larger value or amount of analyte is being measured (23,24).

### 2.10 Statistics

Statistical analyses were conducted in SPSS (version 25). Graphs were made in GraphPad Prism 8. Limit of Detections were calculated using the POD/LOD calculator (25). Maximum log difference was calculated as the upper 95% confidence interval of average of the log difference between all sets of paired samples. Interclass correlation analysis was performed in SPSS on the log-transformed data using a two-way random effects model with average measures, and a type c model with a consistency definition. A two-way random effects model was selected because it models both an effect of operator and the sample, and assumes that both are drawn randomly from larger populations.

## 3.0 RESULTS

### 3.1 THERMOCYCLER COMPARISON

#### 3.1.1 Detection limits of the Open qPCR thermocyclers

The limit of detection 95 (LOD_95_) of the Open qPCR thermocyclers is 63.4 gene copies (GC)/5μL (lower limit 43.7 GC/5μL, upper limit 89.2 GC/5μL, n= 40, based on all qPCR tests). This is approximately 1-log higher than the same assays (Avian schistosomes LOD_95:_ 3.4 GC/5μL;Toxic cyanobacteria LOD_95_: 6.25 GC/5μL ; HF183 LOD_95_: 7.2 GC/5μL) performed using our laboratory ABI 7500/QuantStudio 3 thermocycler. All of these assays have been validated in previous papers, the names, sequences, and the references for the primers and probes are found in Supplementary Table 1. Standard curves performed optimally using the Open qPCR thermocyclers (Table 1).

**Table 1.** Standard curves of each assay performed on the Open qPCR and the core lab machine. Data shown represent the average of 5 runs (each consisting of two internal replicates of each standard). Ideal standard curves have an efficiency of between 0.98 and 0.99, a slope of -3.32, and an efficiency of 100%, which can also be represented as an amplification factor of 2, which suggests product has doubled every cycle.

|  | Copy number in standard | Cycle threshold<br>(average, st dev) |  | r2 | Slope | Amplification<br>factor |
| --- | --- | --- | --- | --- | --- | --- |
| Cyanobacteria mcyE assay |  |  |  |  |  |  |
| Open qPCR | 5000 | 30.1 | 0.27 | 0.980 | -3.05 | 2.100 |
|  | 500 | 33.2 | 1 |  |  |  |
|  | 50 | 36.3 | 1.6 |  |  |  |
| ABI qPCR | 5000 | 28.7 | 1.5 | 0.990 | -3.700 | 1.800 |
|  | 500 | 32.2 | 2.4 |  |  |  |
|  | 50 | 36.1 | 0.91 |  |  |  |
| Human-associated bacteroides (HF183) assay |  |  |  |  |  |  |
| Open qPCR | 5000 | 25.6 | 0.25 | 0.99 | -3.46 | 1.94 |
|  | 500 | 29.2 | 0.24 |  |  |  |
|  | 50 | 32.9 | 0.28 |  |  |  |
| ABI qPCR | 5000 | 24.8 | 0.13 | 0.98 | -4.4 | 1.87 |
|  | 500 | 28.1 | 0.31 |  |  |  |
|  | 50 | 33.7 | 0.26 |  |  |  |
| Pan-avian schistosome assay |  |  |  |  |  |  |
| Open qPCR | 5000 | 27.3 | 0.41 | 0.99 | -3.03 | 2.1 |
|  | 500 | 30.3 | 0.64 |  |  |  |
|  | 50 | 33.3 | 0.52 |  |  |  |
| ABI qPCR | 5000 | 26.9 | 0.52 | 0.99 | -3.03 | 2.1 |
|  | 500 | 30.5 | 0.47 |  |  |  |
|  | 50 | 32.9 | 0.54 |  |  |  |

### 3.2 Comparison between machines

Interclass correlation coefficients (ICC) were calculated to compare CBM partner DNA extracts run on the Chaibio Open qPCR machine, and our laboratory ABI 7500/QuantStudio 3. In 2017, the ICC of the avian schistosomes assay was 0.88 (95% CI: 0.85 lower, 0.90 upper), and in 2018, it was 0.76 (95% CI: 0.56 lower, 0.866 upper), in 2018 this group used 2 Open qPCR machines, and this ICC is a pooled result of both of these machines. In 2018, the ICC of the toxic cyanobacteria assay was 0.57 (95% CI: 0.1 lower, 0.86 upper) (Table 2). Maximum log differences were also calculated and ranged from 1-1.5 depending on the test and year (Table 2).

**Table 2.**
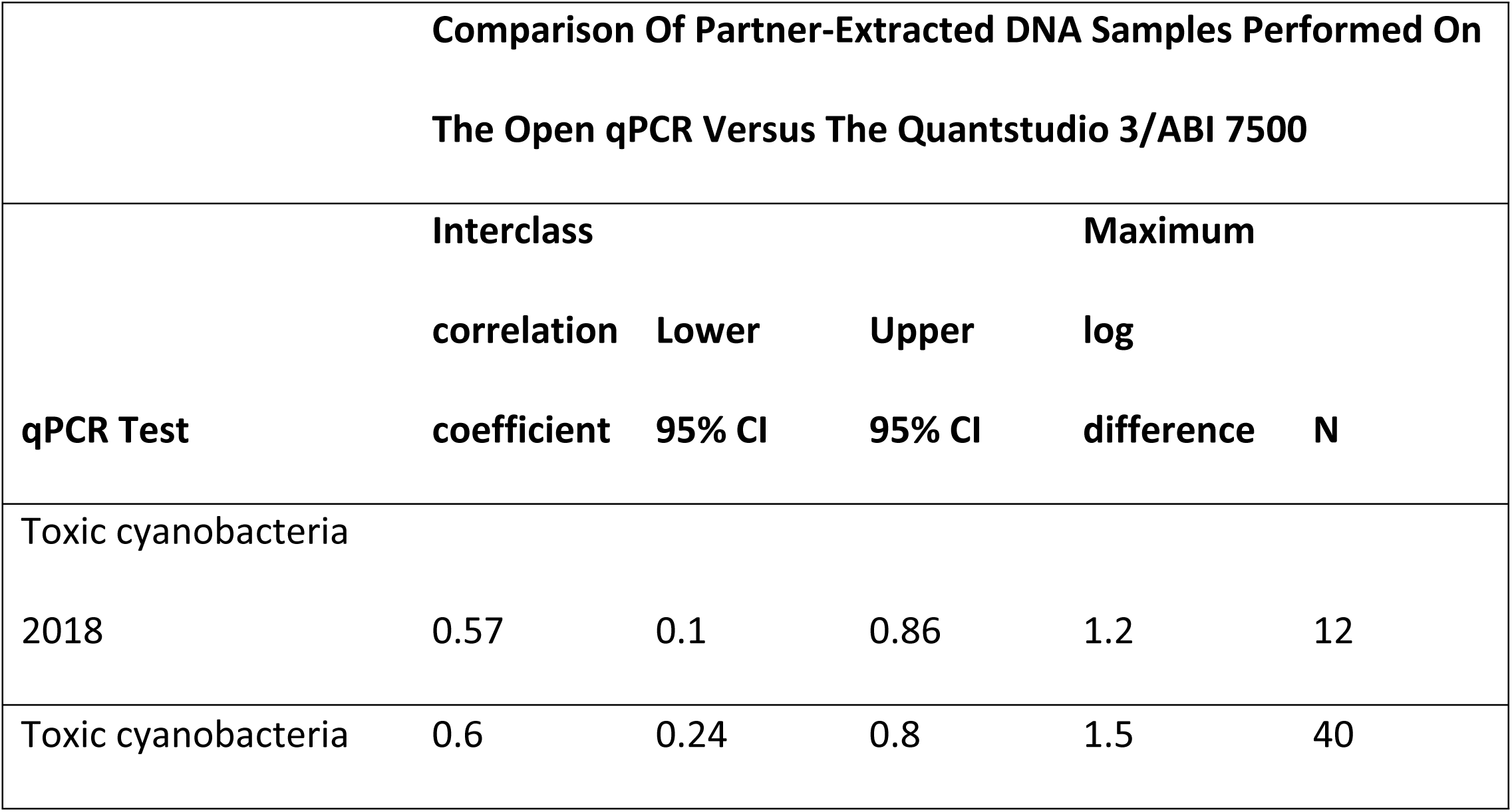
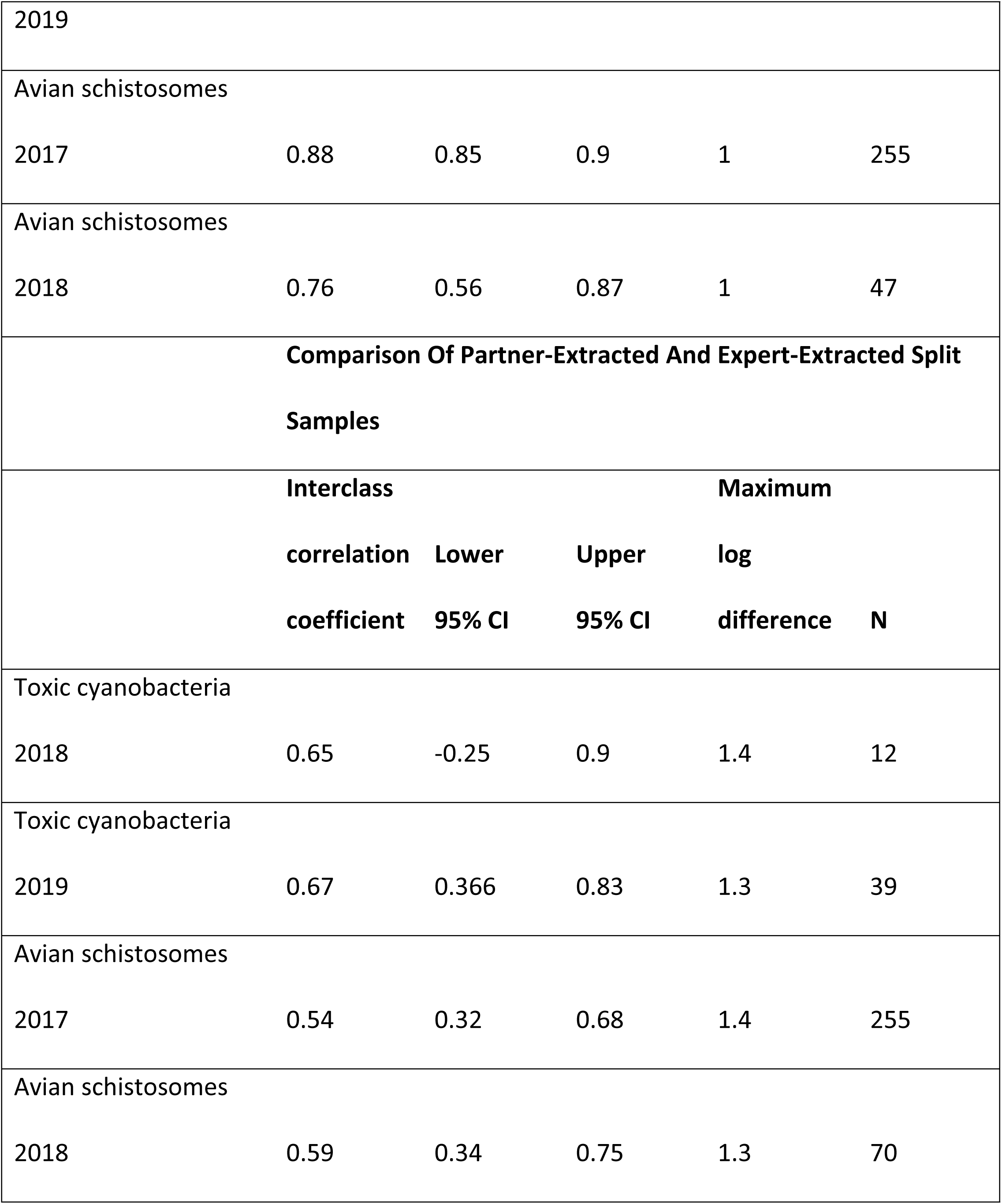
Interclass correlation Coefficients and Maximum Log Difference. comparing the reproducibility of samples run on the Chaibio Open qPCR thermocycler and the ABI 7500 thermocycler/QuantStudio, and

### 3.3 CBM PARTNER COMPARISON

#### 3.3.1 Semi-quantitative analysis using Bland-Altman plots

Reproducibility was assessed using the semi-quantitative Bland-Altman plot. Bland-Altman plots graph the average of two measurements on the X-axis and the difference between these measurements on the Y-axis. The Bland-Altman plot for avian schistosomes monitoring for 2017 and 2018 show a linear pattern at lower copy numbers, but at higher copy numbers show uniform variability (Fig 2). Bland-Altman analysis of the toxic cyanobacteria test shows uniform variability within the limits of agreement (1.96 times the standard deviation). A paired t-test using the log-transformed data was used to compare the within-subject standard deviations of the partner data compared to the lab-generated data. They were significantly different based on an F-test and Welch’s t-test (p < 0.0001, F=6288, mean difference ± SEM: 20326 ± 9843.

**Fig 2.**
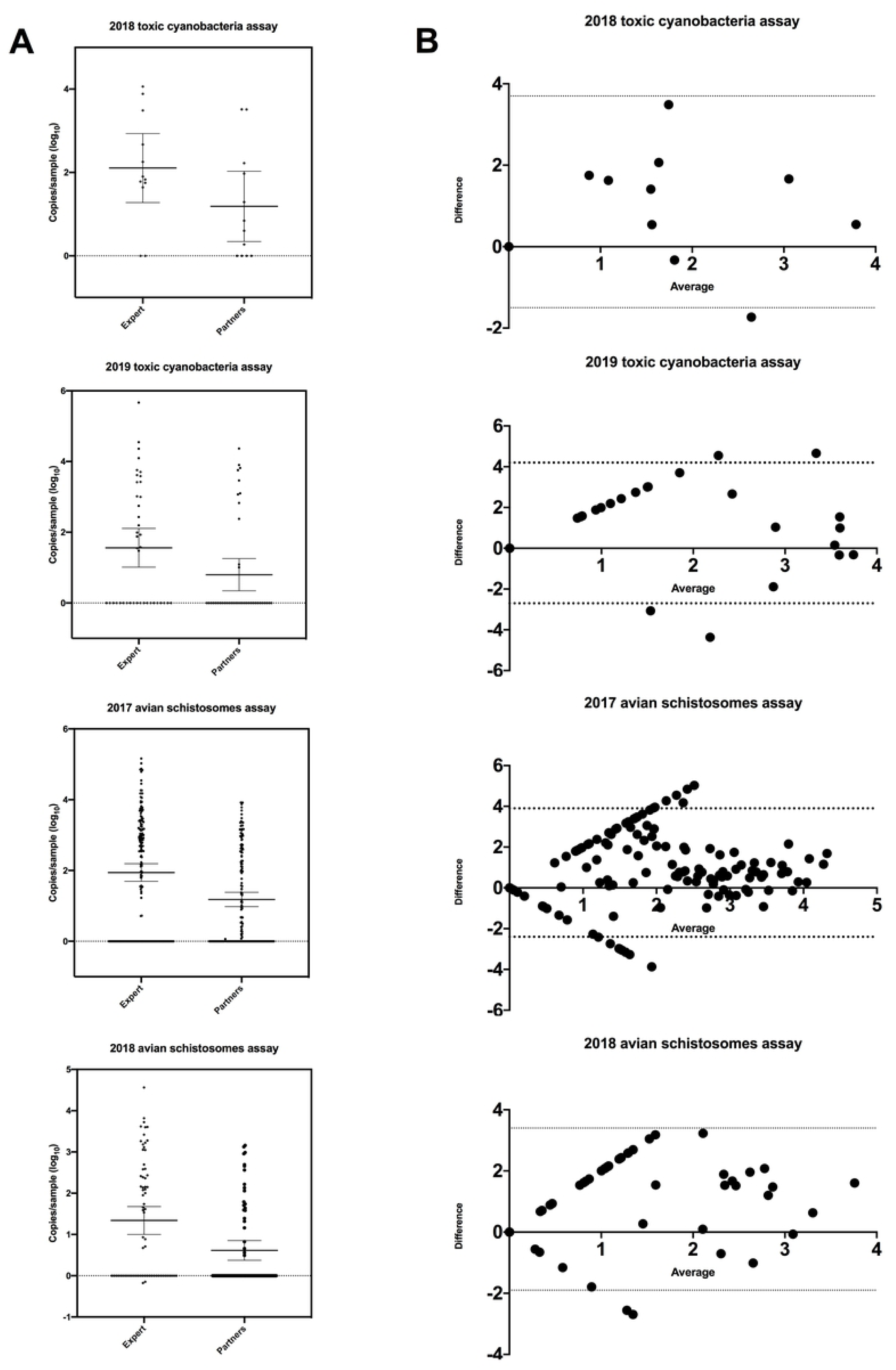
Bland-Altman graphs of the difference between the CBM partners data and the central labs data. Limits of agreement (1.96 times the standard deviation) are bounded by the dotted lines. Top: Agreement of the 2017 Avian schistosome monitoring program. Middle: Agreement of the 2018 avian schistosomes monitoring program. Bottom: Agreement of the microcystin gene monitoring program.

#### 3.3.2 Interclass Correlation Analysis

ICC analysis was performed to compare user and lab samples. In 2017, the Biomeme MI extraction kit was used for swimmer’s itch monitoring. The ICC between user and lab extraction samples was 0.539 (95% CI: 0.320 lower, 0.680 upper). The ICC 2018 for avian schistosomes monitoring was 0.593 (95% CI: 0.344 lower, 0.747 upper). The ICC mcyE was 0.640 (95% CI: -0.250 lower, 0.896 upper) (Table 2). Maximum log differences ranged from 1.3-1.4 (Table 2).

#### 3.3.3 Inhibition controls

PCR Inhibition was tested on partners DNA extractions and on DNA extractions performed in house. Between 5-8% of samples were slightly inhibited in both partner and in house extractions. Cyanobacteria samples were most likely to be inhibited. Inhibited samples were excluded from the analyses in this paper.

### 3.4 QUALITATIVE ANALYSIS

#### 3.4.1 User Perceptions

User perceptions of the program were captured through a written survey that was administered to participants. The questions are available in Supplementary Table 3. Thirty-three percent (33%) of respondents stated that they had some prior knowledge of molecular biology, PCR (polymerase chain reaction), eDNA, or DNA based detection in general prior to the use of the qPCR field method. Fifty percent (50%) reported having low prior knowledge and one participant had no prior knowledge. The same 33% of respondents who reported some knowledge with molecular biology and methods also reported having performed some form of PCR in the past. The rest of respondents reported not having performed PCR (50%) and one participant did not remember. However, prior knowledge did not impact the training all users were provided.

#### 3.4.2 Thematic analysis

User surveys underwent deductive thematic analysis whereby surveys were coded, and then codes were organized into themes (22). The codes identified and relevant excerpts from the surveys are presented in Supplementary Table 4. The first theme identified is “rapidly responding to hazards”. This theme captured the CBM partners’ perceptions on the speed of the qPCR method and their perceived ability to respond to issues quickly. The second theme identified was the question of who controls the CBM monitoring system. This theme emerged from CBM partners expressing a desire for independence and control over the interpretation of results. The third theme identified was that the triangulation of training was valuable in that most CBM partners suggested that the written and video protocols (complemented with a few in person training sessions) were important to them and enhanced their learning. A subtheme that emerged from this theme was “learning and communication”.

## 4.0 DISCUSSION

In this study, the accuracy of a community based qPCR-monitoring system was assessed. We assessed the accuracy of the portable qPCR machines relative to a “core” machine, and the ability for CBM partners to execute the method. Our analyses have demonstrated that a CBM qPCR monitoring program can yield accurate results for different targets (i.e.: eukaryotic versus prokaryotic); however, if the method itself is too time consuming or challenging to be completed by relatively novice CBM partners, a larger scale implementation of the CBM monitoring program could be less reliable.

Our intention was to implement a CBM qPCR system in a real-world context. As Fig 1 details, we began the development of this project by consulting with local stakeholder groups and assessing their interest in the project and what types of biological hazards and surrogates they might be interested in monitoring for. Our goal was to have partners run a sufficient number of tests, not to prescribe a particular test for CBM partners to run. Therefore, we adapted to the needs of our CBM partners and adapted a variety of existing qPCR tests to the field equipment and testing protocol. Additionally, some of the groups we worked with had their own scientific questions they wanted to answer, and we facilitated this.

Our laboratory distributed all materials required to complete testing to users, additionally we prepared all qPCR master mix components (enzyme mix, primers and probes), and aliquoted these into individual reaction tubes for users. The purpose of this was two-fold, to prevent contamination of CBM partners’ qPCR reactions, and for simplicity for partners. Our laboratory facilities are equipped with a PCR clean room, as well as separate pre and post amplification rooms. By preparing reaction tubes and controls, we could prevent CBM partners handling high copy number controls (a likely source of contamination). Additionally, CBM partners were instructed not to open tubes that had undergone qPCR. The Biomeme DNA extraction does not utilize pipettes, but all users were supplied with filter-tipped 20μL pipettes to add their purified DNA into their reaction tubes. Pre-preparing reaction tubes made running qPCR as simple as adding the DNA and pressing “Start” on the Open qPCR machine.

Analysis of the qPCR data was also performed by our laboratory. CBM partners would download their spreadsheets from the Open qPCR and send them either via email or google drive to our labs, where we would analyze control data, and calculate copy numbers and, where possible, organismal numbers for partners. Again, this was done in attempt to preserve the simplicity of the method, and because analysis of qPCR data is complex and requires an expert eye.

The CBM partners participating in this study ran 985 total samples over the two years of this program. Deductive thematic analysis was performed to analyze CBM partner surveys, which is a method of analysis by which codes and theme development were directed by our existing research questions. Three primary themes emerged from this analysis.

The first theme identified was “Rapidly responding to hazards”. Our CBM partners liked that the “time requirement from the qPCR testing method was less than the traditional operational time frame…” However, when asked about the time it took for the method to be completed and if this time was appropriate, all of our CBM partners equated rapidness of the method to a rapid policy response to hazards. This was likely not the reality, as any hazard was dealt with formally through our existing regulatory system which is currently still adapting to the implementation of qPCR methodologies for water monitoring at their core facilities, and for which clear policy frameworks and courses of action do not yet exist for qPCR test results. The only exception to this reality was the avian schistosomes monitoring group, who liked “that [they] could use the next day to change [their] field procedures and experimental designs”.

The second theme was “Independence and verification of a CBM monitoring system”. Two codes that emerged during analysis were CBM partners expressing a desire for more independence and more control over the interpretation of results. Our study was designed to remove data interpretation from participant’s hands, and instead place it in our own hands (with the vision that in a CBM monitoring system that data analysis would be accomplished by a central data processor or the enforcement agency). We thought this would be beneficial because the interpretation of qPCR data is not trivial (especially for quantitative tests that can be correlated to organismal or health outcome levels), and to prevent panic if CBM partners saw positive samples that, while meaningful, might not constitute a real concern. Nonetheless, CBM partners said “the only way these results would be more valuable would be to have a quantitative number which would correlate to specific standard or relative unit conversation chart.” In reality, this was what we were doing for CBM partners, but this group expressed a desire to conduct this independent of our assistance. Additionally, we had a group suggest that they wished the data was published online, “If the data was available or if there was a way to input the data online into a database. Then we could use the results more easily,” they said. Our CBM partners also expressed a desire to validate their results and have access to quality control data. One user suggested “…a visual that compared our results to yours so we have some idea of if we were capturing the results accurately.” Another specifically suggested that, “…third party verification can be one method to enhance validity of the results,” suggesting a desire for some oversight to ensure data quality, but also a desire for CBM partners to know that they are contributing meaningful and accurate results.

One of the biggest challenges for CBM programs is data validation, storage, and visualization. Many communities lack the ability to share CBM data online. However, tools are emerging to address this challenge, including the Lake Observer mobile app through the Global Lake Ecological Observatory Network, the DataStream through the Gordon Foundation, or the ABMI’s NatureLynx. Allowing community partners to upload and visualize their results may help to create a sense that partners are part of something bigger than just their lake. It might allow them to contextualize their results relative to other water bodies, log additional environmental observations, or upload photographs of recreational waters. These apps can also be helpful to track long-term results, or to have the data incorporated into reporting by other agencies.

The third theme we identified was that the triangulation of training was valuable. CBM partners appreciated the three forms of training. Most CBM partners found “the training videos were really useful.” CBM partners found the written protocol useful as a reference, but suggested that after “around 2-3 runs of the machine this resource was no longer needed.” Most CBM partners stressed the importance of the in-person training and one user stated that “the in-person training went a long way in creating and (sic) increased comfort and confidence in the machine.” Studies conducted assessing training in citizen science or CBM projects have found that multiple training sessions can improve data accuracy (26).

A subtheme that emerged during analysis was that CBM partners appreciated the learning process. One user stated that they were “…always up for learning new methodologies to answering scientific questions.” CBM programs are often touted for the positive learning experiences they create, and it is nice to see that ours also had positive learning outcomes for participants (6). A number of CBM partners also suggested that they appreciated the ability to communicate results quickly to their volunteers or to residents on the lakes they worked on. This type of a CBM project could greatly improve science literacy and communication.

The LOD_95_ is the lowest concentration of DNA that can be reliably detected in 95% of samples; it is a measure of sensitivity. The Open qPCR thermocycler has a higher limit of detection when using a Taqman fluorescein probe than our ABI core thermocyclers (63.4 DNA copies/5μL versus >10 DNA copies/5μL across all methods). The field thermocyclers are less sensitive than the core laboratory machine. Understanding this change in detection limit is important to determining if the CBM qPCR system would be effective for a particular test. For instance, if the concentration of the target that might constitute a risk is below the LOD_95_ for the Open qPCR thermocyclers, “risky” samples will appear negative as the thermocycler is not capable of detecting them. For example, when we deployed the human-associated bacteroides HF183 CBM testing for recreational shoreline source tracking in Michigan, USA, our CBM partners reported only a single positive sample. However, when these DNA extracts were analyzed, 22.7% (54/237), were found to be positive for between 15-35 copies DNA/5ul. Seven (0.07%) of these samples approached the LOD_95_ of the Open qPCR thermocycler, and CBM partners detected one of these samples. A recent study found that a HF183 gene copy number of 3220 HF183/100ml exceeds the USEPA benchmark risk of GI illness (10). This level is equivalent to a gene copy number of 161 HF183 GC/5ul—well above the detection limit of the Open qPCR thermocycler. Thus, outbreak scenarios would be clearly discernable. However, this also illustrates an example of how the monitoring project must be clearly rooted in a management outcome. If the intention of the monitoring program is to detect potential outbreak scenarios and initiate action, the increased detection limit is acceptable, yet if the management context is detection of leaking septic areas or source tracking fecal markers on a beach, this detection limit may be inappropriate to answer such questions. This example highlights the importance of working closely with CBM partners to understand their specific monitoring questions, and critically appraising and assessing if CBM qPCR is capable and appropriate to answer these questions.

ICC analysis for the avian trematode assays showed a very high level of agreement between the Open qPCR thermocycler and the core thermocyclers (Table 2). We can expect highly reproducible results between the core machines and the field units. The toxic cyanobacteria test showed much lower levels of agreement between the field thermocycler and the lab thermocycler. We discovered through analyzing the control standards that the heated lid on the field thermocycler was loose, and therefore was failing to engage properly with the tops of the reaction tubes (i.e. machine failure). However, from a quality control perspective, the fact that we were able to detect a probable machine failure with a sample size comparison of merely 11 is extremely promising for future larger scale CBM qPCR systems. It suggests that it would be possible with a relative low number of samples being confirmed by a core facility or quality control partner to detect user or machine error once a baseline level of agreement for a single test had been established.

The comparison between CBM partners performing DNA extraction and myself performing the DNA extraction was first assessed semi-quantitatively using the Bland-Altman plot (Table 2). The results of this analysis for the almost all targets show a linear and negative linear pattern at lower gene copy numbers. This can be due to bias between methods, but can also be caused by a difference in the within-subject standard deviation (24). This seems plausible as users with potentially very different skill levels are performing the two methods. A paired t-test using the log-transformed data was used to compare the within-subject standard deviations. They were significantly different, which suggests that the linear pattern observed is due to an increased variability in CBM partner data.

Partner extracted samples are typically lower in copy number than expert extracted samples (Fig 2). This is likely due to differences in DNA extraction efficiency between the CBM partners and myself. However, it seems more experienced users become better at DNA extraction over time, as both the avian schistosomes monitoring group and the toxic cyanobacteria monitoring groups seem to improve over time. (Fig 2).

Its unsurprised that the ICCs and maximum log differences would be higher when comparing partner and expert extracted DNA samples due to the highly variable nature of DNA extraction, and because the duplicate samples run in the central lab could never be expected to contain exactly the same amount of organism. The ICCs of the DNA extraction comparison ranged from 0.54 to 0.67, with maximum log differences ranging from 1.3 to 1.4 (Table 2). It is important to note that for the avian schistosomes monitoring program, a change was made in 2018 to establish a full functional remote laboratory, and move these partners onto using the Qiagen DNAEasy DNA extraction kit. This change was made at the request of the CBM partners, who would typically collect and analyze hundreds of samples each field season. Details about the equipment in this satellite laboratory can be found in Supplementary Table 2.

Ebentier et al. (2013) conducted a reproducibility analysis of five core laboratories on a panel of microbial source tracking qPCR markers. They calculated reproducibility as the maximum expected log difference (within 95% confidence) between the different laboratories. Their analysis demonstrated reproducibility coefficients for different qPCR assays were highly variable, between 0.09-0.66 log. The methods that were likely to produce higher copy numbers, like Enterococcus qPCR testing via USEPA Method 1611, showed higher reproducibility coeffients than methods that were likely to produce lower copy numbers, like human associated bacteroides marker HF183. They also analyzed the contribution to variability of a variety of factors (the sample itself, equipment, procedures) to the measurement. Their paper concluded that when protocols and reagents were not standardized, agreement between methods decreased. They highlighted the need for standardization of protocols and consumables before implementation of studies involving multi-laboratory experiments (27,28).

The maximum log difference of the CBM qPCR monitoring program higher than the values reported in the Ebentier paper. Reproducibility between the same extract performed by myself and the CBM partners ranged from 0.44 to 1.5 log, and reproducibility coefficients of between partner and expert extracted split samples ranged from 1.3 to 1.4 log (Table 2). It should be noted that the majority of the qPCR methods deployed routinely detected copy numbers in excess of 1 log, thus we might expect higher variability between replicates at these larger copy numbers (Fig 2). CBM qPCR monitoring programs will likely generate data that does have higher variability. It’s important to weigh the pros of a CBM qPCR approach, notably that a CBM qPCR approach may result in increased numbers of samples from across a larger geographic area, and builds relationships and partnerships across sectors.

Rapid monitoring approaches, including CBM qPCR, should be deployed within the context of a policy framework and management response plan that can support acting upon the results generated. The response plan for samples that might constitute a hazard should be clear to CBM partners. If response plans lack transparency, a CBM partner who encounters a sample that contains a high level of an indicator organism, but upon subsequent tests shows low or no risk, might be dismayed by a lack of response by government. A CBM qPCR monitoring system in recreational water would need to prioritize communication and understanding between regulators and CBM partners, and would likely function best when addressing specific objectives(29).

Whether the rapid CBM qPCR monitoring system enables a more rapid response to hazards is yet to be seen; however, CBM qPCR monitoring certainly has the advantage of being able to generate data over a large geographic area and for numerous hazards. It could be adapted to measure organisms not typically considered in monitoring programs; as we have demonstrated in our study, the approach works equally well for eukaryotic hazards like parasitic organisms as it does for the more traditional prokaryotic targets like enteric bacteria. The flexibility inherent in CBM qPCR makes this an attractive and adaptive platform for governments and communities to answer management related questions for their watersheds.

Our vision for the CBM qPCR monitoring system was that data analysis would not occur in the hands of CBM partners (Fig 1). Analysis of qPCR data, while not extremely complex, does require a more comprehensive understanding of qPCR data; additionally, data interpretation is typically the most erroneous component over CBM programs (30,31). Despite our CBM partners desire for independence in data interpretation, we feel that a central ‘expert’ should still be responsible for data interpretation in order to ensure quality in reporting. This could be a single laboratory or a network of QC partners. However, our successes establishing a field laboratory for avian schistosomes and enteric bacteria monitoring in Michigan suggests that a CBM qPCR network could operate effectively within a framework that paired CBM volunteers with quality control partners that could also be operating remotely from the central agency.

Participants in our study expressed a desire to know how well they were performing the method. This highlights an important component of a large-scale CBM monitoring program: a compliance testing system which would test and train potential participants to ensure the method is being conducted appropriately. We believe a this must include third-party verification of a certain percentage of all samples tested. While verification is important to ensure CBM partners are generating reliable results, it is essential that communication be prioritized. This includes responding quickly to results reported by CBM partners when a potential hazard is detected. It also includes being honest with partners about their performance, and willingness by both the CBM and regulatory partners to collect and assess additional samples when clarification or confirmation is required.

## 5.0 Conclusion

To our knowledge, this is the first study to comprehensively test the accuracy of a CBM qPCR water monitoring approach in a real-world context. Our results show that when implemented in a controlled manner, such that a central body controls materials and protocols, results can be highly reproducible. Our study also suggests that CBM partners, whose buy-in would be required for ensuring program longevity, value the method, the data, and what they could do with that data.

CBM qPCR could process a large number of samples from a wide geographical area that could aid beach management for health and invasive species. CBM qPCR could act as a valuable component of an environmental monitoring surveillance system, but could also be a viable option for monitoring and management of rural drinking water systems. qPCR is a platform, and therefore a myriad of diagnostic tests could be deployed as needed in remote locations. While CBM qPCR programs may be more variable than traditional monitoring programs, they could serve as a comprehensive screening system for traditional monitoring programs. In many contexts, CBM qPCR programs could be as accurate as traditional testing and have the potential to replace traditional testing.

## 6.0 ACKNOWLEDGEMENTS

The authors would like to acknowledge the community partners who worked to collect samples and perform qPCR reactions.

## 10.0 SUPPLEMENTARY TABLES

**Table 1S:**
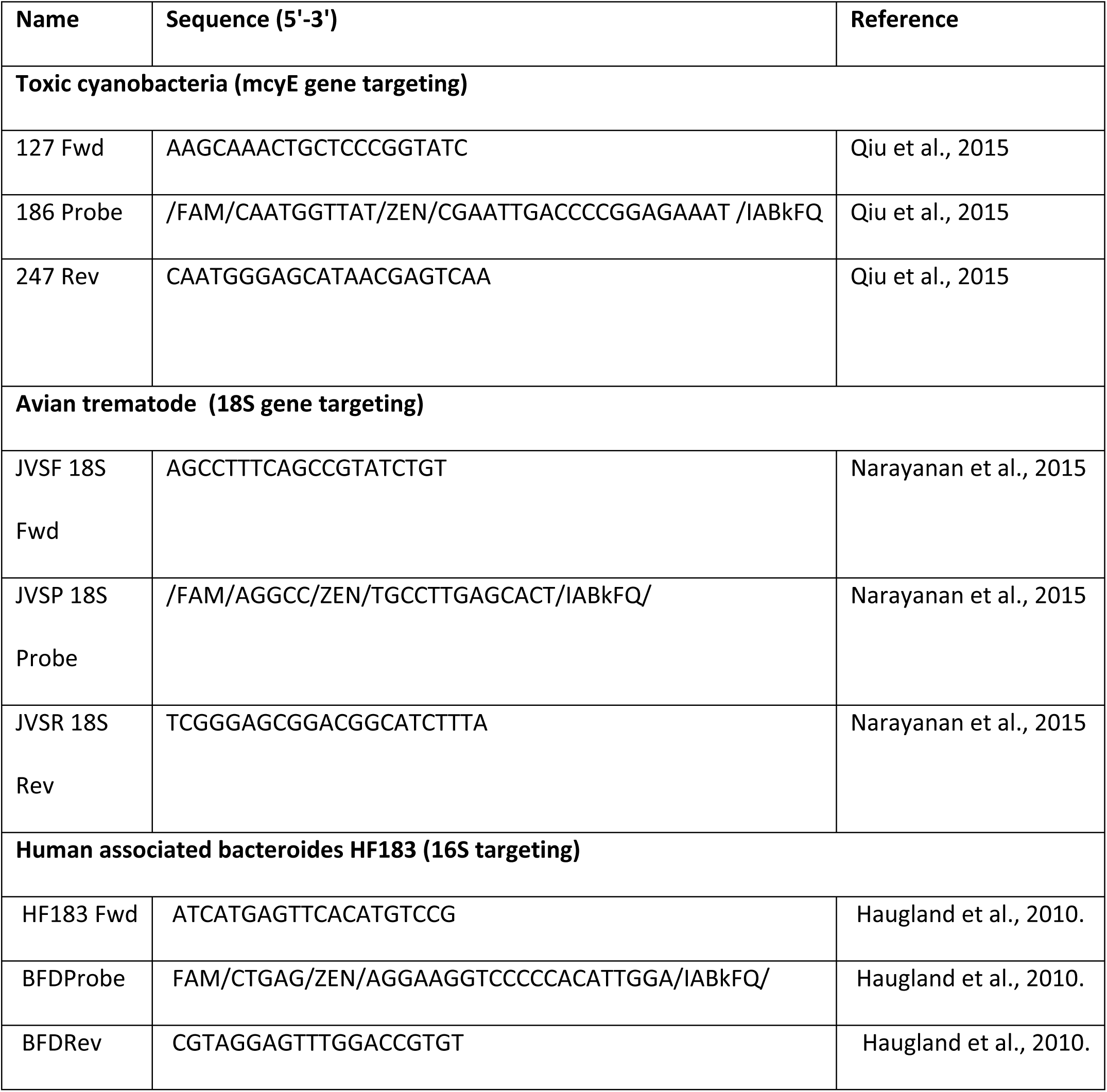
Primers and Probes used in this study.

**Table 2S.**
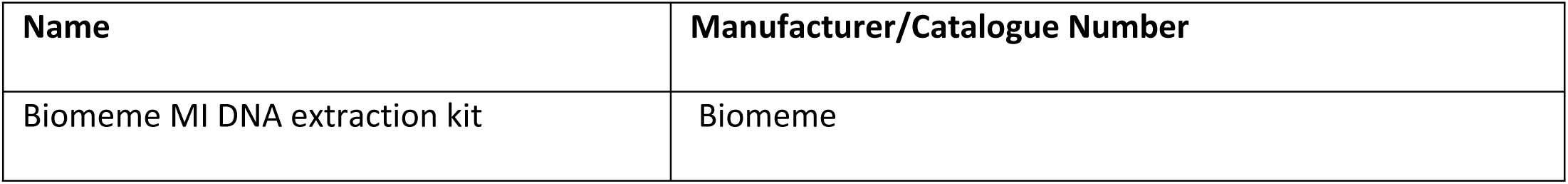
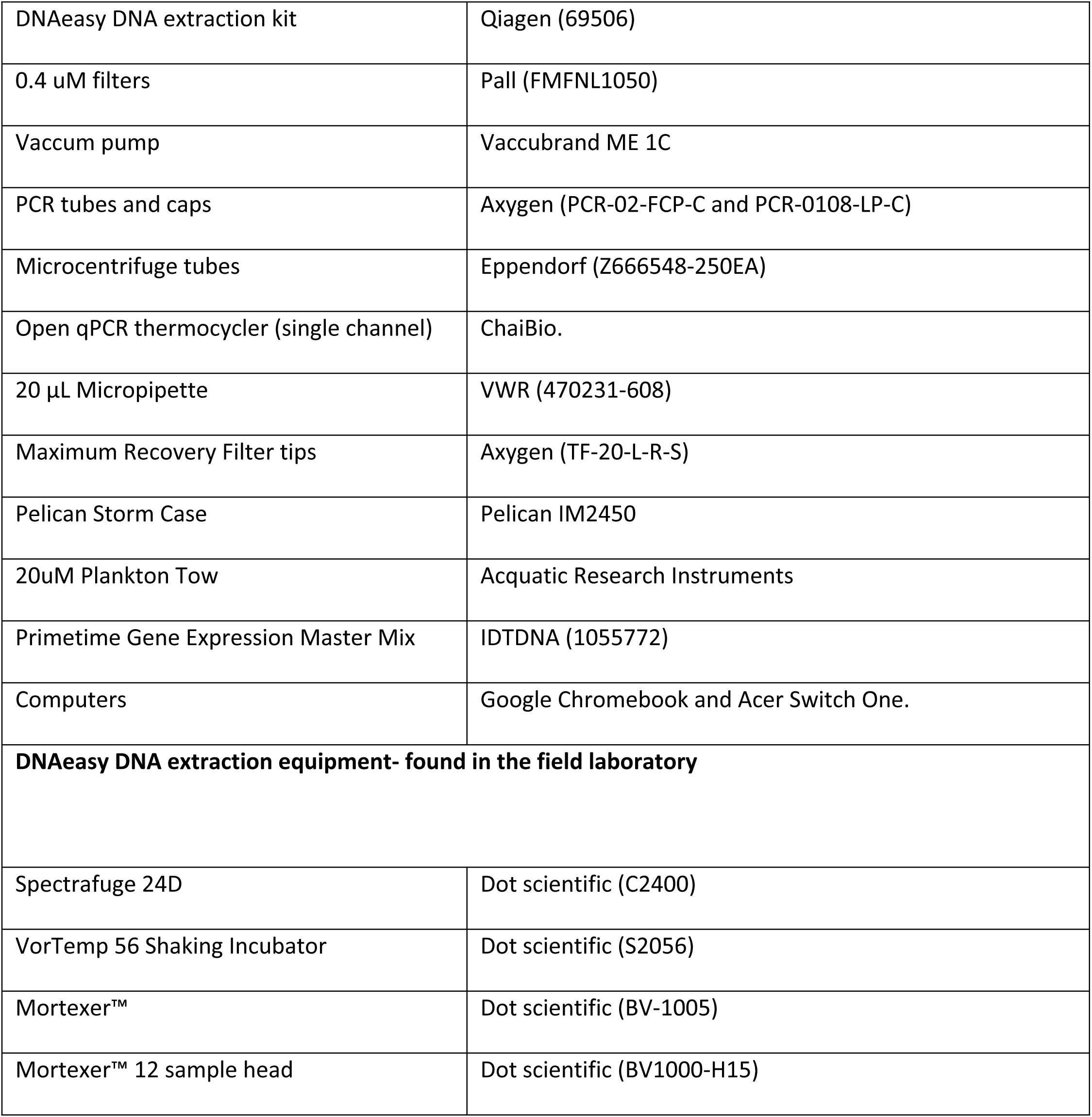
Field method materials used in this study.

**Table 3S.**
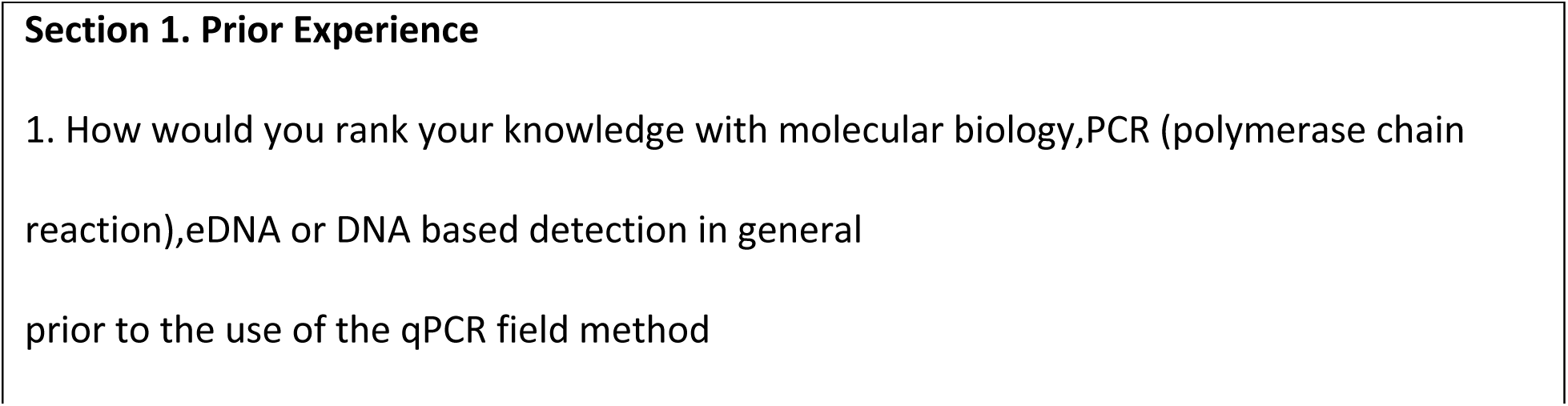
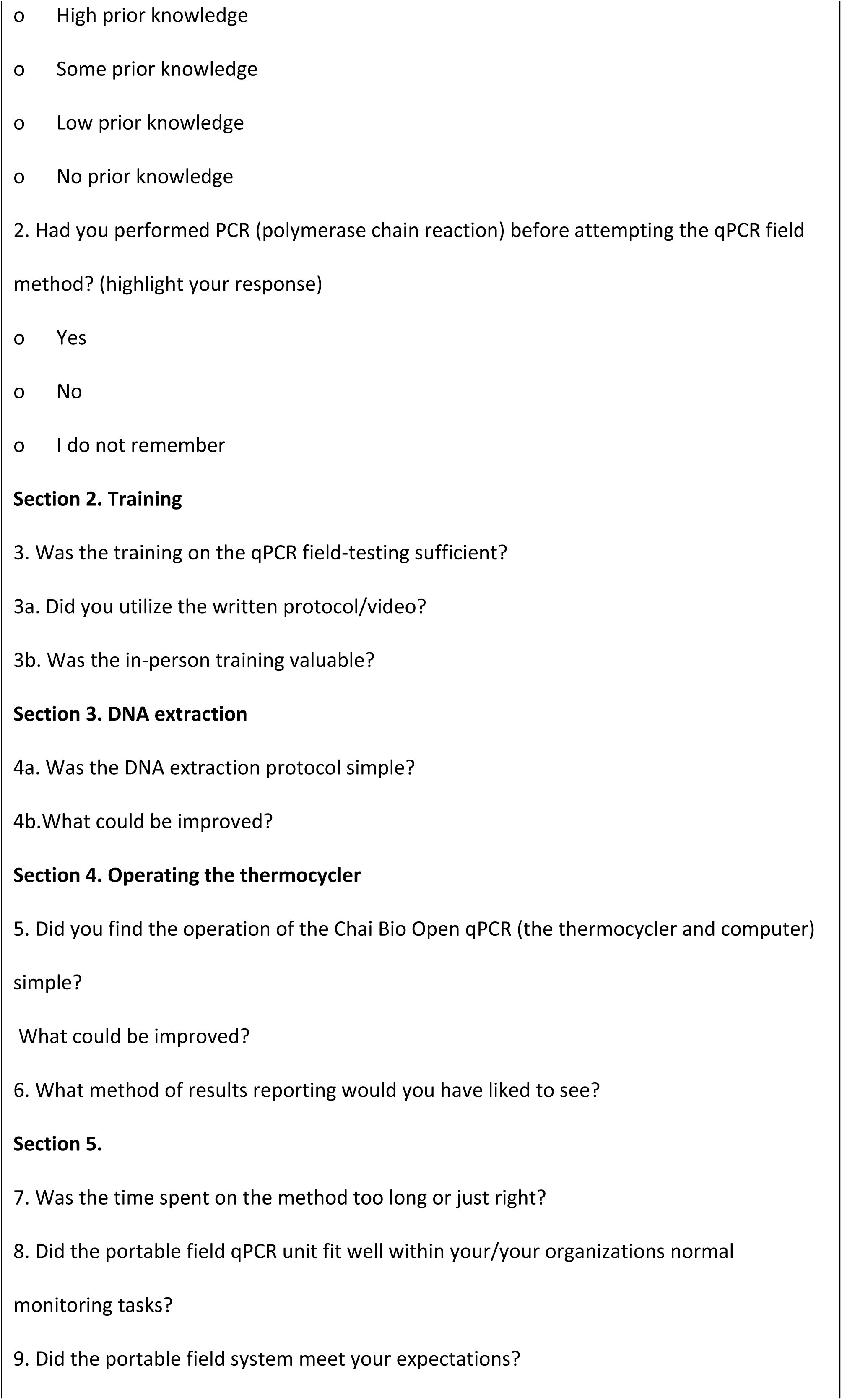
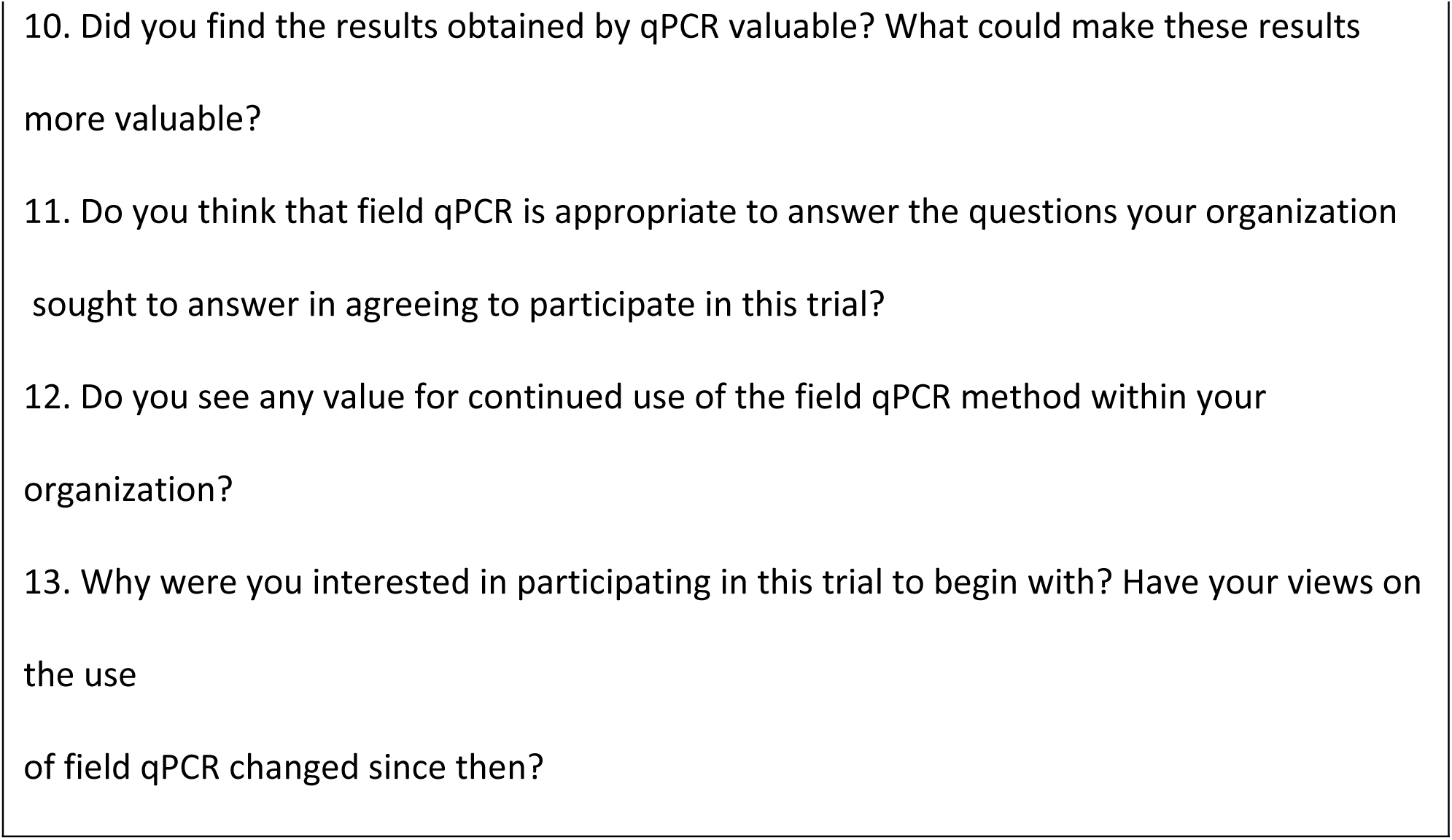
Questions administered to users in survey.

## REFERENCES

1. Conrad CC, Hilchey KG. A review of citizen science and community-based environmental monitoring: issues and opportunities. Env Monit Assess [Internet]. 2011 [cited 2019 Mar 18];176:273–91. Available from: www.eman-rese.

2. Conrad CC, Hilchey KG. A review of citizen science and community-based environmental monitoring: issues and opportunities. Environ Monit Assess [Internet]. 2011 May 17 [cited 2017 Jan 4];176(1–4):273–91. Available from: http://link.springer.com/10.1007/s10661-010-1582-5

3. Obama Administration. Accelerating Citizen Science and Crowdsourcing to Address Societal and Scientific Challenges. The White House. 2015.

4. Serrano F, Holocher-Ertl T, Kieslinger B, Sanz F, Silva C. White Paper On Citizen Science For Europe. Annual Review of CyberTherapy and Telemedicine. 2013.

5. Lakshminarayanan S. Using Citizens to Do Science Versus Citizens as Scientists. Ecol Soc [Internet]. 2007 Dec 28 [cited 2019 Mar 18];12(2):resp2. Available from: http://www.ecologyandsociety.org/vol12/iss2/resp2/

6. Trumbull DJ, Bonney R, Bascom D, Cabral A. Thinking scientifically during participation in a citizen-science project. Sci Educ [Internet]. 2000 Mar 1 [cited 2017 Oct 4];84(2):265–75. Available from: http://doi.wiley.com/10.1002/%28SICI%291098-237X%28200003%2984%3A2%3C265%3A%3AAID-SCE7%3E3.0.CO%3B2-5

7. Whitelaw G, Vaughan H, Craig B, Atkinson D. Establishing the Canadian community monitoring network. Environ Monit Assess [Internet]. 2003 [cited 2019 Mar 18];88(1–3):409–18. Available from: https://link.springer.com/content/pdf/10.1023/A:1025545813057.pdf

8. Wade TJ, Calderon RL, Sams E, Beach M, Brenner KP, Williams AH, et al. Rapidly measured indicators of recreational water quality are predictive of swimming-associated gastrointestinal illness. Environ Health Perspect [Internet]. 2006 Jan [cited 2019 May 3];114(1):24–8. Available from: http://www.ncbi.nlm.nih.gov/pubmed/16393653

9. Wymer LJ, Brenner KP, Martinson JW, Stutts WR, Schaub SA, Dufour AP. The EMPACT Beaches Project : Monitoring in Recreational Waters. Environ Prot. 2005;

10. Boehm AB, Soller JA, Shanks OC. Human-Associated Fecal Quantitative Polymerase Chain Reaction Measurements and Simulated Risk of Gastrointestinal Illness in Recreational Waters Contaminated with Raw Sewage. Environ Sci Technol Lett. 2015;

11. Cao Y, Sivaganesan M, Kelty CA, Wang D, Boehm AB, Griffith JF, et al. A human fecal contamination score for ranking recreational sites using the HF183/BacR287 quantitative real-time PCR method. 2018 [cited 2019 May 6]; Available from: https://doi.org/10.1016/j.watres.2017.10.071

12. Ibelings BW, Backer LC, Kardinaal WEA, Chorus I. Current approaches to cyanotoxin risk assessment and risk management around the globe. Harmful Algae [Internet]. 2014 Dec [cited 2016 Oct 4];40:63–74. Available from: http://www.ncbi.nlm.nih.gov/pubmed/26435706

13. Abou Tayoun AN, Burchard PR, Malik I, Scherer A, Tsongalis GJ. Democratizing Molecular Diagnostics for the Developing World. Am J Clin Pathol [Internet]. 2014 Jan 1 [cited 2017 Sep 26];141(1):17–24. Available from: https://academic.oup.com/ajcp/article-lookup/doi/10.1309/AJCPA1L4KPXBJNPG

14. Wood CS, Thomas MR, Budd J, Mashamba-Thompson TP, Herbst K, Pillay D, et al. Taking connected mobile-health diagnostics of infectious diseases to the field. Nature. 2019.

15. Baum F, MacDougall C, Smith D. Participatory Action Research. J Epidemiol Community Health [Internet]. 2006 Oct [cited 2019 Jun 7];60(10):854–7. Available from: http://www.ncbi.nlm.nih.gov/pubmed/16973531

16. Blankespoor HD, Reimink RL. The Control of Swimmer’s Itch in Michigan: Past, Present, and Future. MICHIGAN Acad XXIV. 1991;7–23.

17. Rudko SP, Reimink RL, Froelich K, Gordy MA, Blankespoor CL, Hanington PC. Use of qPCR-Based Cercariometry to Assess Swimmer’s Itch in Recreational Lakes. EcoHealth. 2018;

18. Qiu Y, Yuan T, Zurawell R, Huang Y. Rapid Detection and Quantitation of Microcystin-Producing Microcystis Using Real-Time PCR. J Mol Biomark Diagn [Internet]. 2013 [cited 2017 May 30];S5. Available from: http://dx.doi.org/10.4172/2155-9929.S5-006

19. Sipari H, Rantala-Ylinen A, Jokela J, Oksanen I, Sivonen K. Development of a chip assay and quantitative PCR for detecting microcystin synthetase E gene expression. Appl Environ Microbiol [Internet]. 2010 Jun [cited 2016 Jul 28];76(12):3797–805. Available from: http://www.ncbi.nlm.nih.gov/pubmed/20400558

20. Rudko SP, Ruecker NJ, Ashbolt NJ, Neumann NF, Hanington PC. Investigating Enterobius vermicularis as a novel surrogate of helminth ova presence in tertiary wastewater treatment plants. Appl Environ Microbiol [Internet]. 2017 Mar 24 [cited 2017 Apr 10];AEM.00547-17. Available from: http://www.ncbi.nlm.nih.gov/pubmed/28341675

21. Fereday J, Muir-Cochrane E. Demonstrating Rigor Using Thematic Analysis: A Hybrid Approach of Inductive and Deductive Coding and Theme Development. Int J Qual Methods [Internet]. 2006 Mar 29 [cited 2019 Dec 16];5(1):80–92. Available from: http://journals.sagepub.com/doi/10.1177/160940690600500107

22. Braun V, Clarke V. Using thematic analysis in psychology. Qual Res Psychol [Internet]. 2006 Jan [cited 2019 May 6];3(2):77–101. Available from: http://www.tandfonline.com/doi/abs/10.1191/1478088706qp063oa

23. Martin Bland J, Altman DG. Stastical Methods for Assessing Agreement Between Two Methods of Clinical Measurement. Lancet. 1986;

24. Bartlett JW, Frost C. Reliability, repeatability and reproducibility: analysis of measurement errors in continuous variables. Ultrasound Obstet Gynecol [Internet]. 2008 Apr 1 [cited 2019 Mar 20];31(4):466–75. Available from: http://doi.wiley.com/10.1002/uog.5256

25. Wilrich C, Wilrich P-T. Estimation of the POD Function and the LOD of a Qualitative Microbiological Measurement Method. [cited 2015 Jun 22]; Available from: http://www.ingentaconnect.com/content/aoac/jaoac/2009/00000092/00000006/art00020

26. Ratnieks FLW, Schrell F, Sheppard RC, Brown E, Bristow OE, Garbuzov M. Data reliability in citizen science: learning curve and the effects of training method, volunteer background and experience on identification accuracy of insects visiting ivy flowers. Methods Ecol Evol [Internet]. 2016 [cited 2019 May 3];7(10):1226–35. Available from: http://cygnus-biologystudentjournal.wikispaces.

27. Ebentier DL, Hanley KT, Cao Y, Badgley BD, Boehm AB, Ervin JS, et al. Evaluation of the repeatability and reproducibility of a suite of qPCR-based microbial source tracking methods. Water Res [Internet]. 2013 Nov [cited 2017 Jul 20];47(18):6839–48. Available from: http://linkinghub.elsevier.com/retrieve/pii/S0043135413005484

28. Shanks OC, Sivaganesan M, Peed L, Kelty CA, Blackwood AD, Greene MR, et al. Interlaboratory comparison of real-time pcr protocols for quantification of general fecal indicator bacteria. Environ Sci Technol. 2012;

29. Barzyk TM, Huang H, Williams R, Kaufman A, Essoka J. Advice and Frequently Asked Questions (FAQs) for Citizen-Science Environmental Health Assessments. Int J Environ Res Public Health [Internet]. 2018 [cited 2019 Jun 7];15(5). Available from: http://www.ncbi.nlm.nih.gov/pubmed/29751612

30. Hunter J, Alabri A, van Ingen C. Assessing the quality and trustworthiness of citizen science data. Concurr Comput Pract Exp [Internet]. 2013 Feb 1 [cited 2017 Dec 15];25(4):454–66. Available from: http://doi.wiley.com/10.1002/cpe.2923

31. Cohn JP. Citizen Science : Can Volunteers Do Real Research ? Bioscience. 2008;58(3):192–7.

